# Solving the stochastic dynamics of population growth

**DOI:** 10.1101/2022.11.15.516663

**Authors:** Loïc Marrec, Claudia Bank, Thibault Bertrand

**Affiliations:** Institut für Ökologie und Evolution, Universität Bern, Baltzerstrasse 6, CH-3012 Bern, Switzerland; Swiss Institute of Bioinformatics, 1015 Lausanne, Switzerland; Department of Mathematics, Imperial College London, South Kensington Campus, London SW7 2AZ, United Kingdom

## Abstract

Population growth is a fundamental process in ecology, evolution, and epidemiology. The population size dynamics during growth are often described by deterministic equations derived from kinetic models. Here, we simulate several population growth models and compare the size averaged over many stochastic realizations with the deterministic predictions. We show that these deterministic equations are generically bad predictors of the average stochastic population dynamics. Specifically, deterministic predictions overestimate the simulated population sizes, especially those of populations starting with a small number of individuals. Describing population growth as a stochastic birth process, we prove that the discrepancy between deterministic predictions and simulated data is due to unclosed-moment dynamics. In other words, the deterministic approach does not take into account the variability of birth times, which is particularly important at small population sizes. We evaluate different moment-closure approximations and show that they do not satisfactorily reduce the error between analytical predictions and simulated data. We present two novel solutions to the stochastic growth dynamics, one of which applies to any population growth model. We show that our solution exactly quantifies the dynamics of a community composed of different strains and correctly predicts the fixation probability of a strain in a serial dilution experiment. Our work sets the foundations for a more faithful modeling of community dynamics. It provides tools for a more accurate analysis of experimental results, including the inference of important growth parameters.

## 1. INTRODUCTION

Population growth is at the heart of fundamental processes in cell biology, evolution, ecology, and epidemiology, from the expansion of bacteria colonies and large-scale animal populations to the spread of an infectious disease or the propagation of an advantageous mutation. Predicting population growth dynamics is thus paramount in ecology and controlling the spread of infections. Indeed, in the context of epidemiological modeling, understanding the transiently increasing and steady-state size of the infected population can help us devise strategies to control population growth and mitigate its spread. The outspoken goal of population modeling is to accurately describe the variation in the number of individuals in a population.

Historically, deterministic models are most commonly used to describe population dynamics [1–3]. In these models, the population size is generically described by a continuous variable whose temporal dynamics are governed by an ordinary differential equation. Whereas most of these models are nonlinear— which means that analytical progress is not impossible but limited in some cases [4]—it is often relatively simple and computationally fast to obtain accurate numerical solutions, possibly explaining their widespread use. In theoretical ecology, a paradigmatic model of population growth is the well-known logistic equation whose study traces back to as early as the middle of the nineteenth century [5]. The logistic differential equation was initially derived from introducing a self-limiting property in the growth of a biological population to the unconstrained Malthusian exponential growth model [6] and was rediscovered independently later on [7–9]. The derivation of Verhulst’s logistic growth model stemmed from the observation that unhindered exponential population growth is unrealistic because even in the absence of predation relations, intraspecies competition for environmental resources such as food or habitat will lead to a characteristic saturation level, an upper bound on the population size known as the *carrying capacity*. Owing to its ease of use, the simplest logistic growth was used to model biological systems at all scales from the population growth of micro-organisms [10, 11] to that of large mammal herds [12] and fish schools [13].

Further refinements to the logistic growth function led to the development of a *generalized logistic growth model* [4], which captures several commonly used population growth models including Blumberg [14], Richards [15] and Gompertz [16] growth models. Whereas amenable to easy progress and qualitative predictions, these deterministic models are not entirely faithful to the growth of a real population which is inherently stochastic [17, 18]. This stochasticity results from both intrinsic (e.g., demography) and extrinsic (e.g., environmental change) noise [19, 20]. More recent studies have shown that deterministic and stochastic approaches yield critically different results [21–24]. Although it was long considered that the dynamics of a large volume stochastic system could be well described by deterministic equations [25], it is now clear that this criterion alone is not sufficient [26]. The range of validity of deterministic models is put in question. Even if new conditions for a deterministic equation to describe well the stochastic dynamics of a population have been outlined [26], they are not exhaustive and quantitative methods to overcome this discrepancy are missing. Recognizing these limitations, stochastic models have proved helpful in epidemiology and ecology for the past decades [27–36].

Many studies use deterministic equations to fit experimental or empirical data and estimate essential biological and environmental parameters. Logistic growth models have recently been used in microbiology [37, 38] as well as in epidemiology, e.g., to model the Covid-19 pandemic [39, 40]. In the latter example, several studies used logistic-like equations to estimate the basic reproductive number of a virus during an outbreak [41–44], whereas others used compartmental models such as SEIR [45–47]. In both cases, a prediction based on deterministic models carries the risk of poorly estimating parameters of interest, such as the reproductive number, crucial to implementing political measures to slow down the disease’s spread. Identifying when a deterministic equation does not describe well the average dynamics of stochastic population growth, understanding the reasons for this disagreement, and proposing solutions to remedy it, is thus of paramount importance.

In this work, we leap forward by solving the stochastic dynamics of population growth in the absence of deaths analytically. This resolution allows us to identify the extent to which a deterministic approach is a good approximation of the growth dynamics and to lay the foundation for future inference methods of growth parameters. We consider several classical population growth models. First, we model the population growth as a stochastic birth process and simulate stochastic realizations of these kinetics. We compare their ensemble average to the predictions of the respective deterministic models. We show that the deterministic approach generically overestimates the average population size; this error in prediction is shown to be larger when the initial number of individuals is very low. To explain the reason behind this discrepancy, we derive a master equation formalism describing the stochastic population growth dynamics and the moment equations. We find that the difference between the population size estimated by the deterministic equation and the mean of the simulated data is due to unclosed moment dynamics. We show that some moment-closure approximations greatly reduce the difference, which remains globally significant. Instead, we derive an exact solution to the stochastic population growth. Finally, we apply our results and show that our solution leads to a better prediction of the dynamics of two competing strains and the probability of fixation of a mutant in a serial passage experiment.

## 2. BIAS OF DETERMINISTIC APPROACHES

### 2.1. Pure-birth models

Given the inherent stochasticity of population growth processes, we first establish whether a deterministic equation correctly describes the mean trajectory of stochastic growth. We consider four distinct growth models belonging to the *generalized logistic growth models*: Blumberg, Gompertz, Logistic, and Richards models [4]. Our choice was motivated by their widespread use to fit experimental or empirical data to estimate growth parameters in microbiology and ecological communities [38, 48, 49]. These kinetic models differ by their *per capita* growth rates, *b*_*N*_.

Under Malthusian growth, the *per capita* growth rate is constant and independent of the population size *N* ; we denote as *b* this intrinsic birth rate—also called *exponential birth rate* or *Malthusian parameter*. Note that simple unbounded exponential growth would occur if no restrictions were imposed on the population size (e.g., nutrients, available space). In the growth models considered here, the *per capita* growth rate is explicitly dependent on the population size. To model environmental constraints such as availability of space or food, one then generically introduces a carrying capacity *K* that limits the population size *N* to the range *N*_0_ ≤ *N* ≤ *K* assuming no deaths, where *N*_0_ is the initial population size. Specifically, as the population size increases, the *per capita* birth rate *b*_*N*_ decreases to vanish when *N* = *K*.

For a general Logistic growth model, the deterministic equation describing the dynamics of *N* reads

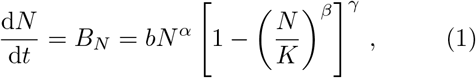

where *B*_*N*_ denotes the population growth rate. For simplicity, we consider the case *α* = 1, but our results stand in the more general case. The population growth rates for each of the four nonlinear models considered here are provided in table 1 along with those of the exponential growth model.

**Table 1.** Population growth models. List of specific population growth models used with their associated exponents and population birth rates *B*_*N*_. These are derived from the *generalized logistic growth model* introduced in equation (1). The exponents *α, β* and *γ* allow tuning the symmetry, maximum and inflection of the population growth rate *B*_*N*_ (see figure 1). Birth rate, population size, and carrying capacity are denoted by *b, N*, and *K*, respectively.

| Exponential | Logistic | Blumberg | Richards | Gompertz* |
| --- | --- | --- | --- | --- |
| $\alpha = 1$ | $\alpha = 1$ | $\alpha$ | $\alpha = 1$ | $\alpha = 1$ |
| $\beta = 0$ | $\beta = 1$ | $\beta = 1$ | $\beta$ | $\beta \rightarrow 0$ |
| $\gamma = 0$ | $\gamma = 1$ | $\gamma$ | $\gamma = 1$ | $\gamma = 1$ |
| $B_N = bN$ | $B_N = bN(1 - N/K)$ | $B_N = bN^\alpha(1 - N/K)^\gamma$ | $B_N = bN(1 - (N/K)^\beta)$ | $B_N = bN \log(K/N)$ |

\* The generalized logistic growth model converges to the Gompertz model when the *per capita* growth rate is divided by $\beta$ and the limit $\beta \rightarrow 0$ is taken (see [4] for details).

As seen in figure 1, the four population growth models chosen here display very different birth rate curves. Birth rates in all models are non-monotonic and vanish when *N* → 0 and *N* → *K* by construction. In other words, these models impose that a population of size zero cannot grow, and no population can grow beyond the carrying capacity. We note that the Logistic model displays a symmetry around the population size *N* = *K/*2, whereas in both the Blumberg and Richards models, the exponents *β* and *γ*, respectively, offer an extra degree of freedom to tune the shape of the growth rate curve and in particular, its asymmetry. We note that the population size at inflection, i.e., when the population growth rate is maximum, is given by

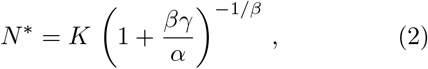

and is thus dependent on exponents *β* and *γ* for the Blumberg and Richards models, respectively. The Gompertz model shows the fastest growth of all at small population sizes. As shown in figure 2(a), the population size in all deterministic models follows a sigmoid curve—also called an S-shape curve—with its characteristic initial phase with slow growth, exponential growth phase, finally followed by a stabilization of the population size at a finite steady-state population size

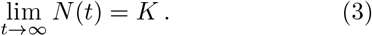

For the Logistic, Richards, and Gompertz models, full analytical solutions are available [4]. These are given by

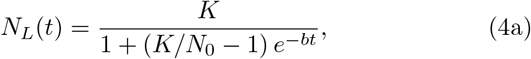

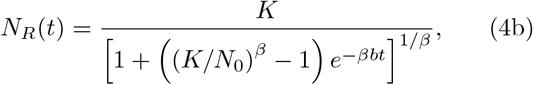

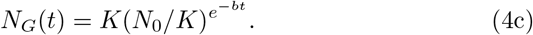

Those are represented in figure 2(a). Note that no analytical (i.e., closed-form) solution exists for the Blumberg model; in this last case, we proceeded with a direct numerical solution of equation (1).

**Figure 1.**
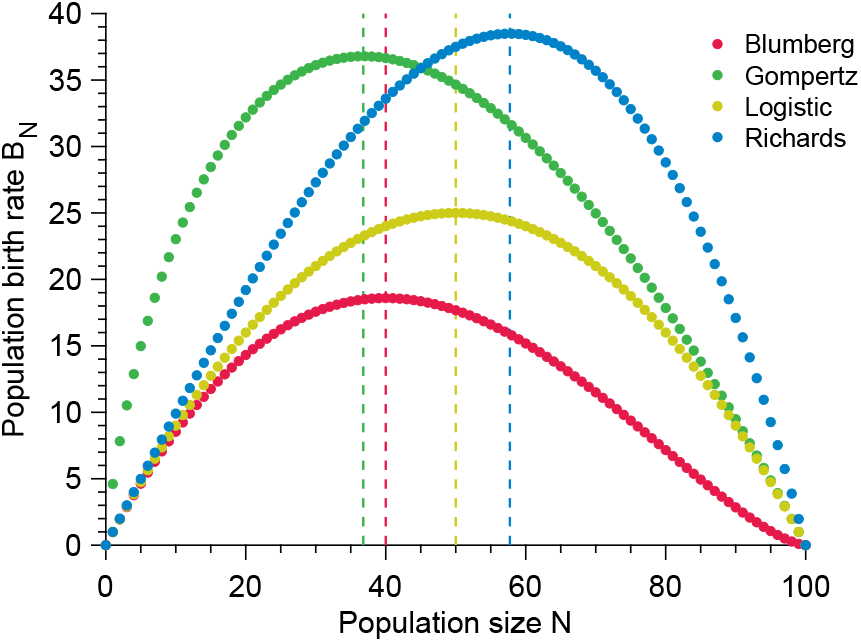
Different growth models display different population birth rates. Vertical dashed lines show the location of the optimal birth rate for each model. Parameter values: carrying capacity *K* = 100, birth rate *b* = 1, exponents *α* = 1, *β* = 2 and *γ* = 1.5.

**Figure 2.**
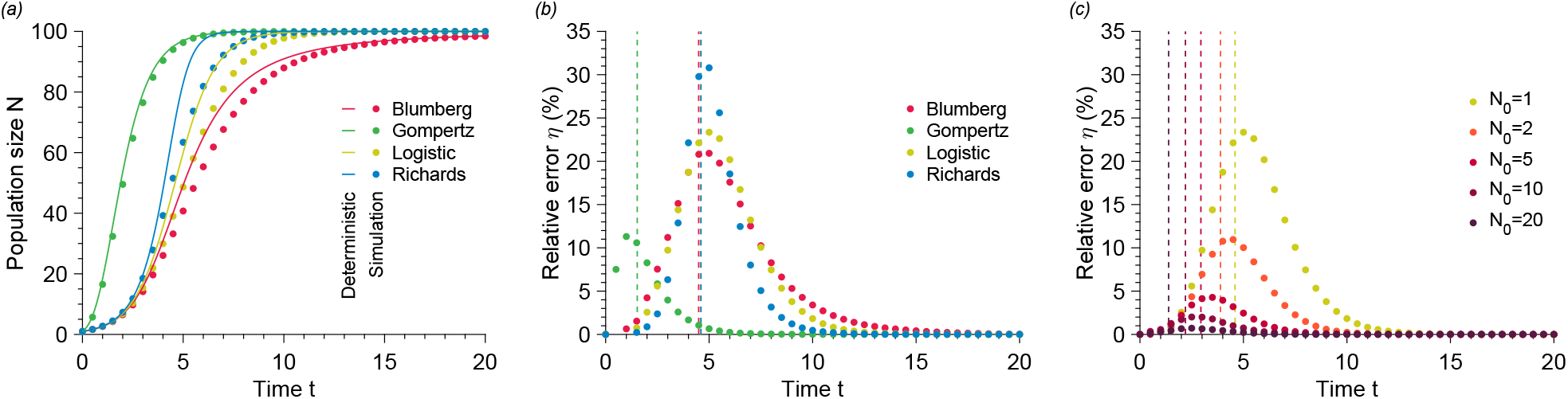
Solutions obtained from deterministic models fail to predict the mean stochastic population size. (a) Population size *N* versus time *t* for different growth models. The data points show simulated data averaged over 10^4^ stochastic realizations. The solid lines correspond to the numerical resolution of the deterministic models (see equation (1)). (b) Relative error *η* versus time *t* for different growth models calculated using data from (a). The vertical dashed lines represent the inflection times at which d^2^*N/*d*t*^2^ = 0 or equivalently when the population growth rate reaches a maximum. (c) Relative error *η* versus time *t* for different initial population sizes *N*_0_ for the Logistic model. The data points show simulated data averaged over 10^5^ stochastic realizations. The vertical dashed lines represent the inflection times. Parameter values: carrying capacity *K* = 100, initial population size *N*_0_ = 1 (in (a) and (b)), birth rate *b* = 1, exponents *α* = 1, *β* = 2 and *γ* = 1.5.

However, as pointed out earlier, population growth is inherently stochastic. We, therefore, evaluated the validity range of the above deterministic descriptions by simulating individual stochastic trajectories for the four growth models introduced above using a Gillespie algorithm [50, 51] [see electronic supplementary material (ESM)]. To this end, we recast our problem into a pure-birth process for which population growth results from an individual *A* reproducing at size-dependent rate *b*_*N*_ following the elementary reaction

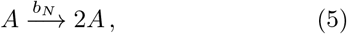

with *B*_*N*_ = *b*_*N*_ × *N*. Naturally, multiple stochastic models may lead to the same deterministic model under a mean-field approximation. Here, we focus on one particular microscopic scenario. Still, as we will argue in the next section, other formulations, including those where birth and death processes are taken into account explicitly, lead to even more drastic disagreement. We average all our results over 10^5^ independent stochastic trajectories to obtain the time-dependent average population size.

As shown in figure 2(a), we observe a significant difference between the deterministic predictions and the stochastic mean population size starting from a single individual. To quantify this disagreement, we calculated the relative error *η* defined as

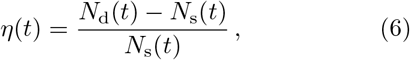

where *N*_d_(*t*) and *N*_s_(*t*) are respectively the timedependent population sizes predicted by the deterministic model and measured in our stochastic simulations. We observed relative errors as large as 30% independent of the carrying capacity (see ESM, figure S2). Figure 2(b) shows that the largest error for the parameters chosen is obtained for the Richards model, whereas the smallest is for the Gompertz model (still at a substantial max(*η*) ≈ 12%). We note that the quantitative value of the relative errors measured for the Blumberg and Richards models depend on the choice of exponents *β* and *γ*, respectively. Nevertheless, the measured errors remain significant over a wide range of exponents (see ESM, figure S1). For all the growth models studied here, the deterministic dynamics given by equation (1) overestimates the population size at all times. It is interesting to note that the maximum error is located around the inflection point (i.e., at *t* such that d^2^*N/*d*t*^2^ = 0) predicted by the deterministic equation; we note that this corresponds to the population size where the population birth rate reaches its maximum and starts decreasing.

Furthermore, we observe that the error uniformly decreases as *N*_0_, the initial number of individuals in the population, increases (see figure 2(c)). This discrepancy limits the range of validity of the deterministic models as the initial number of individuals in the population is assumed to be small in many applications, e.g., patient zero in a disease spreading scenario, single cell mutation in a mutation fixation experiment, small number of cells in a bacterial colony expansion, etc. To summarize, the deterministic equation fails to describe the average stochastic trajectory. Although the relative error depends on the specific growth model, and thus on the *per capita* birth rate, it remains significant in all cases tested and increases with decreasing initial population size. Importantly, the relative error between deterministic predictions and measured mean stochastic population size is independent of the carrying capacity. Thus the discrepancy does not vanish in the limit of large population sizes as is commonly assumed (see ESM, figure S2).

Stochastic population growth is a Poissonian jump process. Therefore, the times between jumps from size *N* to *N* + 1 are exponentially distributed with parameter *B*_*N*_. For small initial population sizes *N*_0_, the rates at which the population initially grows are low (see figure 1), e.g., the rate of the first reproduction is given by *bN*_0_ [1 − (*N*_0_ */K*)^*β*^]^λ^ for the generalized logistic model. In turn, this implies that early in the process, the distributions of reproduction times are heavy-tailed, leading to a large variance in the population size consistent with observations from singlecell experiments [52–56]. We postulate that this large variance accumulated over the growth process is responsible for the disagreement between the deterministic and the mean stochastic trajectories; this is in particular consistent with our observations that: (1) the relative error increases when the initial population size decreases and (2) the relative error is maximal around the inflection point where the exponential distribution of reproduction times is the tightest. Interestingly, when *K* → ∞ (large volume limit), the rate of first reproduction converges to *bN*_0_ and so is entirely controlled by the initial population size confirming that the observed disagreement remains valid in this limit. Finally, we note that the above deterministic models are mean-field models which intrinsically assume an underlying population size distribution peaked around its mean in contrast to the wide population size distributions observed in the stochastic models.

### 2.2. Birth-death processes

In the previous section, we focused on populations that can only increase in size over time. This assumption, which entails neglecting the death of individuals, is predominant in microbiology, where the models used to fit population growth data do not explicitly include death rates [37, 38, 57]. Similarly, pharmacodynamic models, which aim at quantifying how antibiotics inhibit growth or kill cells, commonly replace the birth rate with a net birth rate (i.e., birth rate minus death rate) in equation (1) [58, 59]. In this way, the population grows if the net birth rate is positive, decreases if it is negative, and remains constant for zero net growth rates. However, stochastic population growth can also be modeled as a Poissonian Markov jump process where births and deaths are distinct events leading to distinct changes in population size: *N* → *N* + 1 and *N* → *N* − 1, respectively.

We do not expect deterministic models to be fair better in the presence of explicit death events with small initial population sizes. For the sake of simplicity and without loss of generality, we add to equation (1) a linear death term leading to the modified differential equation d*N/*d*t* = *bN*^*α*^ [1 − (*N/K*)^β^]^λ^ −*dN*.

We further simulate a stochastic birth-death process known to lead to this deterministic equation in the mean-field limit using once again a Gillespie algorithm [50, 51]. Not only do we expect a deterministic approach to fail to describe the dynamics of the average population growth over a large number of stochastic realizations as in the death-free case but we show that strikingly it also fails to predict the correct steadystate population size.

Indeed, figure 3(a) shows that deterministic models do not predict quantitatively either time-dependent population sizes or steady-state population sizes averaged over many stochastic realizations. We argue that this difference is due to rapid extinctions, which are frequent occurrences when considering low initial population sizes and large ratios of death rate to birth rate [see figures 3(b) and (c)]. Although early extinctions are more likely when the death rate is higher than the birth rate, demographic stochasticity leads to extinction with probability 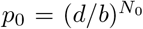 (or 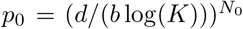 for the Gompertz model), which is non-zero *in all cases*. These early extinction events are not taken into account in deterministic approaches.

**Figure 3.**
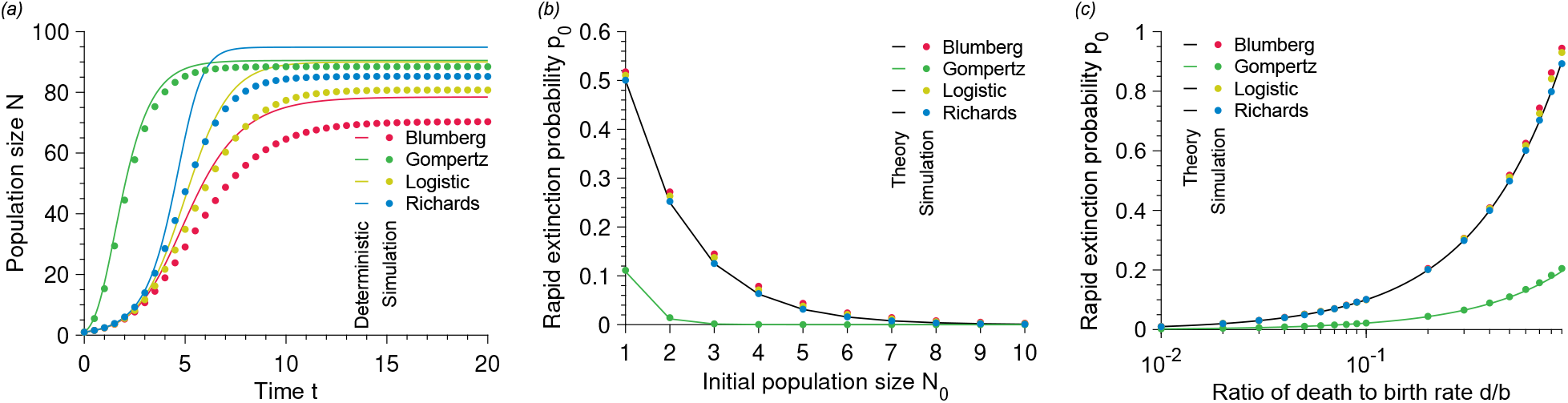
Deterministic approach performs worse with non-zero death rates. (a) Population size *N* versus time *t* for different population growth models. (b) Rapid extinction probability *p*_0_ versus initial population size *N*_0_ for different population growth models. (c) Rapid extinction probability *p*_0_ versus ratio of death to birth rate *d/b* for different population growth models. In every panel, the solid lines the deterministic predictions, and each point results from simulated data averaged over 10^5^ stochastic realizations. Parameter values: carrying capacity *K* = 100, birth rate *b* = 1, death rate *d* = 0.1 (in (a)) and 0.5 (in (b)), initial population size *N*_0_ = 1 (in (a) and (c)).

In summary, we conclude that deterministic formalism is not a good predictor of the average population growth dynamics, even for large carrying capacities in the presence or the absence of explicit death events. We also note that the discrepancy between deterministic and average stochastic population sizes worsens as the initial population size decreases. We further conclude that deterministic approaches fail at predicting the steady-state population size when death events are explicitly introduced. In the following, we focus on the pure-birth process, which is already of great interest in microbiology, as we pointed out earlier. In the next section, we adopt a stochastic approach to describe the population growth and obtain an exact analytical solution for the population size distribution at all times.

## 3. ERROR REDUCTION BY MOMENT-CLOSURE APPROXIMATIONS

To identify the reasons behind the poor performance of the deterministic equation, we return to a stochastic formalism. Generically, any population growth in the absence of death may be described by a stochastic birth process whose rates are defined by the underlying deterministic model [66] (see table 1 for examples). Let us consider a population whose number of individuals at time *t* is denoted by *N*, whereas its initial population size is *N*_0_. As in equation (5), we consider that each individual in the population replicates with the same *per capita* rate *b*_*N*_. Here, the population size increases from *N* to *N* + 1 individuals at a total rate *B*_*N*_, where *B*_*N*_ was defined in table 1 for several population growth models.

We focus on finite-sized populations that grow in a constant environment with a carrying capacity *K*. To fully account for the stochasticity inherent to demographic noise, we use a microscopic and probabilistic description in continuous time of the birth events within the population. More specifically, we write a system of differential equations describing the probabilities 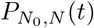 that a population has a given size *N* at a given time *t* knowing that it started with *N*_0_ individuals. Because the growth rates vanish when *N* → *K*, the size of our population is at most *K*, with the state *N* = *K* being an absorbing state, i.e., the population indefinitely remains in this state once reaching it for the first time. Put simply, our stochastic process, while continuous in time, has a finite discrete number of possible states. Here, we assume that the population jumps between successive sizes with a rate dependent on its current size leading to a fully coupled system of equations.

This system of differential equations, formally called the *master equation*, governs the time evolution of the probabilities 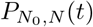. Writing the master equation for a stochastic jump process requires one to think about gain and loss terms to the probabilities 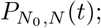; for our system, it reads [67, 68]

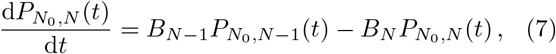

where the first term on the right-hand side of equation (7) is a gain term corresponding to an increase in the population size from *N* − 1 to *N* individuals via a birth event, whereas the second term is a loss term corresponding to the population size transitioning from *N* to *N* + 1 individuals. Although writing down the master equation for a stochastic jump process is often easy, computing the formal solution of master equations is arduous and has been an active field of investigation for decades [67–69].

Importantly, the master equation (7) contains all information about the growth dynamics; in particular, as it governs the probability distribution 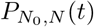, it contains all information to compute the averaged population size trajectory over time. Rather than solving directly equation (7), we derive an equation governing the moments of 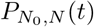, where the *m*-th moment of the population size is defined as

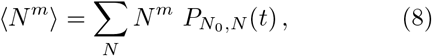

with *m* a positive integer. For instance, the equation governing the time-evolution of the first moment (i.e., *m* = 1), which corresponds to the mean population size, reads

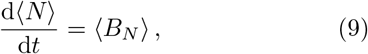

whereas the equation for the second moment (i.e., *m* = 2) satisfies

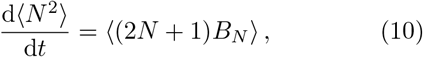

where again averages are defined as 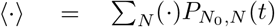.

The moment equations are closed for linear population birth rates as in the Malthusian (or exponential) growth model. Taking for the sake of simplicity and without loss of generality *b* = 1, the population birth rate of the exponential model is given by *B*_*N*_ = *N*. Writing out the moment equations, we obtain, for instance,

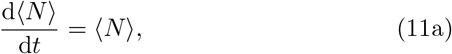

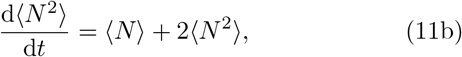

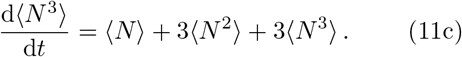

First, the average population size given by equation (11a) shows that the deterministic model accurately describes the average stochastic population size dynamics. Secondly, the equation for the second moment (11b) is closed as it only depends on the second moment itself and the first moment, which can be obtained analytically by solving equation (11a). Similarly, the third moment (11c) is closed as well; the equation for the *m*^th^ moment only depends on the *m* first moments, and so by solving the moment equations sequentially, we obtain a closed equation for any moment of the distribution. This way, important distribution properties such as its variance or skewness can be studied analytically.

Meanwhile, the moment dynamics are unclosed for nonlinear population birth rates, which is often the case for finite populations. In other words, the equation for *m*^th^ moment may involve higher-order moments, leading to an infinite hierarchy of moment equations that is not usually solvable. Approximation techniques to get around this problem exist, and here, we discuss the accuracy of the most commonly used.

The most basic way to get around this problem is to apply a so-called *mean-field approximation*. The mean-field approximation relies on the crucial assumption that the distribution of populations sizes is well-peaked so that 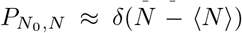, where *δ* is the Kronecker-delta function. This approximation naturally leads to the approximations ⟨*N*⟩ ≈ *N*, and ⟨*B*_*N*_⟩ ≈ *B*_*N*_, which enable us to recover the deterministic limit given by equation (1). More formally, a Kramers-Moyal expansion in combination with a diffusion approximation of the master equation (7) leads to the same resulting deterministic equation [67] (see ESM for details). As we argued earlier, this deterministic model has been very popular, for instance, in population genetics, as it simplifies calculations and circumvents the need for the master equation framework [70–73]. Importantly, we argued in section 2 that the distributions of population sizes were wide (see also figure 5). So unsurprisingly, the mean-field approximation fails to describe satisfactorily the population size averaged over stochastic realizations.

Going beyond the mean-field approximation requires us to close the hierarchy of moment equations; these methods are called *moment-closure approximations*. They have been extensively used to provide analytic approximations to nonlinear stochastic population growth models [64, 65, 74]. In the following, we focus on the Logistic growth model and proceed to several common moment-closure approximations. Writing the first few moment equations for the Logistic growth model, we obtain

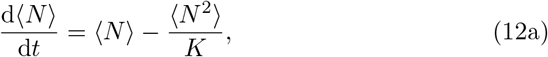

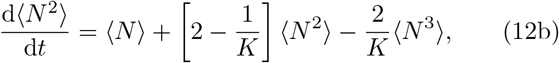

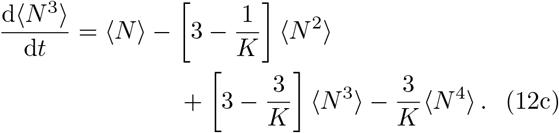

Notably, the equation for the first moment ⟨*N*⟩ depends on the second moment *N* ⟨^2^⟩, whereas the equation for the second moment ⟨*N* ^2^⟩ depends on the third moment ⟨*N* ^3^⟩, and so on. To close this hierarchy of moment equations, two routes are often employed: (i) closure methods can rely on a cumulant truncation procedure, in which the *k* first cumulant equations are approximated by setting all cumulants of order higher than *k* to 0 [32, 63], (ii) closure methods can also be based on assumptions on the form of the underlying distribution of population sizes 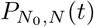 [27, 31, 62, 64, 74]. Most recently, the latter method has been extensively used; in these latter approximations, one often focuses only on the first two moments.

Here, we test several common moment-closure approximations that express ⟨*N* ^3^⟩ as a function of the first two moments ⟨*N*⟩ and ⟨*N* ^2^⟩, allowing us to close the first two moment equations (12a) and (12b). We report all moment-closure approximations tested here in table 2. As shown in figure 4, all moment-closure approximations tested here show a significant disagreement with the simulated stochastic mean population. Whereas the Binomial, separable derivative-matching (SDM), and mean-field approximations overestimate the mean population size, the New-Poisson, Nåsell-Poisson and Normal momentclosure approximations underestimate it. We report absolute relative errors ranging from ∼ 6% for the Binomial approximation to ∼ 26% for the normal approximation. Indeed, classical moment-closure approximations fail in problems with very skewed underlying probability distributions, for which accurate knowledge of the higher-order moments is needed [74]. Figure 5 shows that our population growth dynamics are plagued by large skewness. In other words, figure 5 shows that the population size can have many different values at a given time during stochastic growth.

**Table 2.** Moment-closure approximations. Common moment-closure approximations where the third moment ⟨ *N* ^3^ ⟩ is expressed as a function of the first moment ⟨ *N* ⟩ and the second moment ⟨ *N* ^2^ ⟩.

| Moment-closure approximation | Third moment $\langle N^3 \rangle$ |
| --- | --- |
| Binomial [60] | $2(\langle N^2 \rangle - \langle N \rangle^2)^2 / \langle N \rangle - \langle N^2 \rangle + \langle N \rangle^2 + 3\langle N^2 \rangle \langle N \rangle - 2\langle N \rangle^3$ |
| Lognormal [34, 61] | $\langle N^2 \rangle^3 / \langle N \rangle^3$ |
| Nåsell-Poisson [60] | $\langle N \rangle + 3\langle N^2 \rangle \langle N \rangle - 2\langle N \rangle^3$ |
| New-Poisson [60] | $\langle N^2 \rangle - \langle N \rangle^2 + 3\langle N^2 \rangle \langle N \rangle - 2\langle N \rangle^3$ |
| Normal [62–64] | $3\langle N^2 \rangle \langle N \rangle - 2\langle N \rangle^3$ |
| Separable Derivative-Matching (SDM) [65] | $\langle N^2 \rangle^3 / \langle N \rangle^3$ |

**Figure 4.**
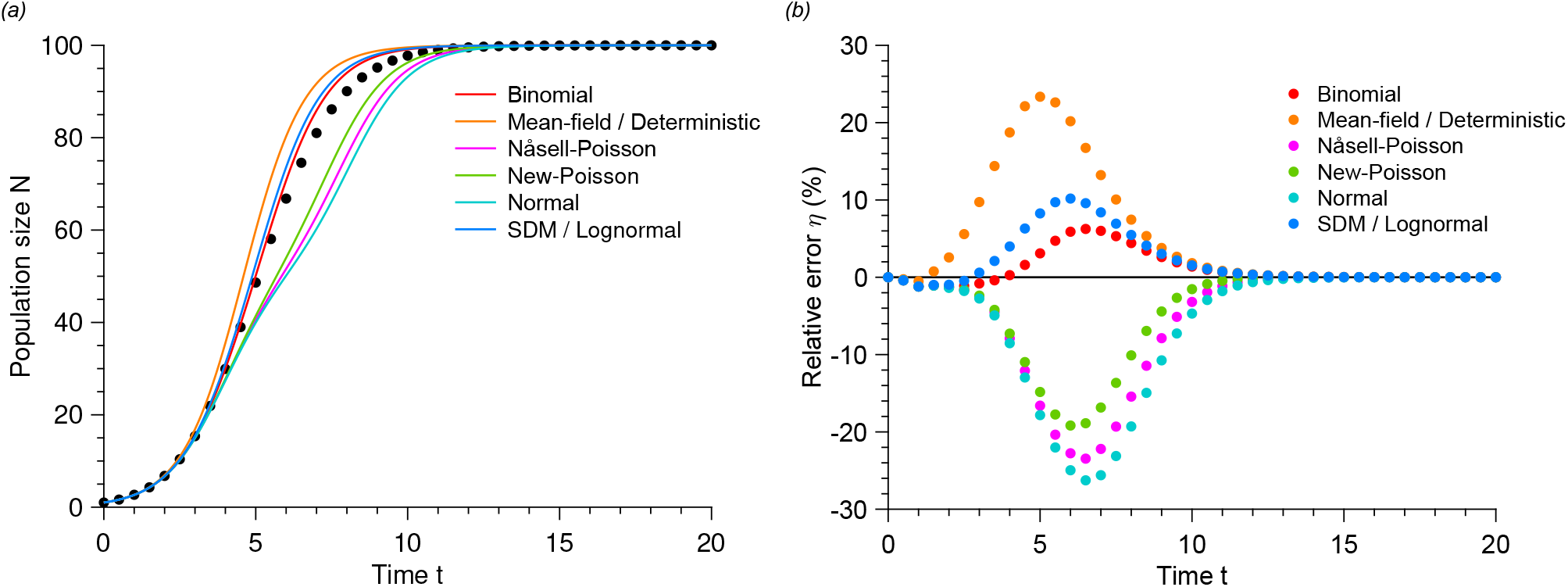
Moment-closure approximations do not satisfactorily reduce the error. (a) Population size *N* versus time *t* for moment-closure approximations with the Logistic model. The data points show simulated data averaged over 10^5^ stochastic realizations. The solid lines correspond to the moment-closure approximations (see Table 2). (b) Relative error *η* versus time *t* for different moment-closure approximations. *η* is calculated using data from Panel (a). Parameter values: carrying capacity *K* = 100, initial population size *N*_0_ = 1 and birth rate *b* = 1.

**Figure 5.**
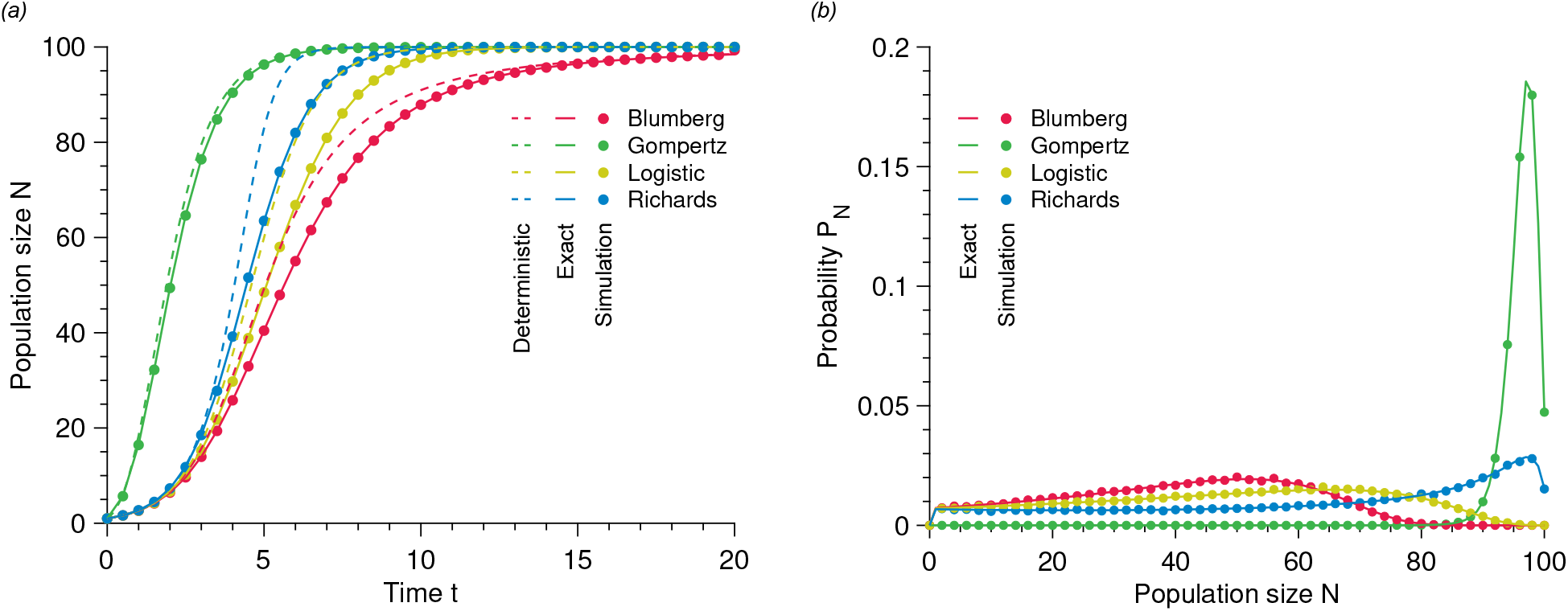
Exact solution to the nonlinear population growth problem. (a) Population size *N* versus time *t* for different population growth models. The data points show simulated data averaged over 10^5^ stochastic realizations. The solid lines correspond to our solution, whereas the dashed lines represent the deterministic equation. (b) Probability *P*_*N*_ of having *N* individuals at *t* = 5. Parameter values: carrying capacity *K* = 100, initial population size *N*_0_ = 1, birth rate *b* = 1, exponents *α* = 1, *β* = 2 and *γ* = 1.5.

While these moment-closure approximations generally reduce the error in predicting the dynamics of population growth, we note that they cannot generically be applied to the other growth models considered in this paper. Indeed, the right-hand side of equation (9) depends on the terms ⟨(1 − *N/K*)^*γ*^*N*⟩, ⟨log(*N*)*N*⟩ and ⟨*N*^*β*+1^⟩ for the Blumberg, Gompertz, and Richards models, respectively. These are not easily expressed in terms of combinations of moments of the distribution. Since these nonlinear population growth models are widely used, it is crucial to devise a generically applicable method. However, we recognize that moment-closure approximations allow growth dynamics to be described from one or two ordinary differential equations, which may be easier than a fully probabilistic description.

## 4. TOWARDS AN EXACT SOLUTION TO STOCHASTIC POPULATION GROWTH

### 4.1. First approach: transition rate matrix

A first approach to try and solve the master equation directly is to recast it in the language of Markov chains. Namely, stochastic population growth can be interpreted as a continuous-time, *K*-state Markov jump process with transition rate matrix **R** = (*R*_*ij*_)_1≤*i*≤*K*,1≤*j*≤*K*_, where the matrix element *R*_*ij*_ with *i* ≠ *j* denotes the rate at which the population switches from value *N* = *i* to *N* = *j*. The diagonal elements of the transition rate matrix are generically fixed by enforcing conservation of total probability, which implies ∑_*i*_ *R*_*ij*_ = 0, so that *R*_*jj*_ = − ∑_*i*≠*j*_ *R*_*ij*_.

As the population sizes potentially vary from 1 to *K*, we first define **P**(*t*) as the column vector of dimension *K*

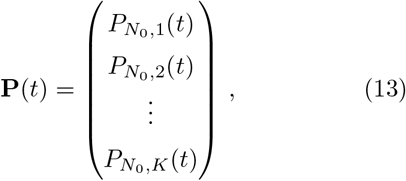

with the *i*^th^-component corresponding to the time-dependent probability of having a population of size *i*. We then rewrite equation (7) in matrix form as

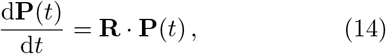

where the *K* × *K* transition rate matrix **R** is defined as

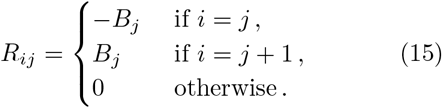

The solution to equation (14) can generically be written as a weighted superposition of the *K* eigenvectors **v**_*k*_ of the transition rate matrix multiplied by an exponential function with rate given by the associated eigenvalue *α*_*k*_. Namely, we write

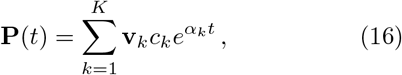

a vector whose components give us the time-dependent probability that the population size is *N* given that it was *N*_0_ at *t* = 0,

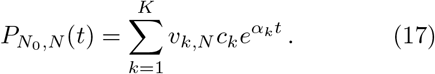

Note that the coefficients *c*_*k*_ are obtained by the imposition of the initial conditions and here must satisfy 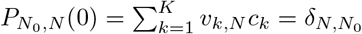, where *δ* is the Kronecker delta. Furthermore, *c*_*k*_ = 0 for 1 ≤ *N < N*_0_ since a population size lower than *N*_0_ cannot be reached. That is because death events are not considered here, so the population size can only increase over time.

As the transition rate matrix **R** is lower triangular, the eigenvalues are equal to the diagonal entries of the matrix, and we obtain *α*_*k*_ = − *B*_*k*_, 1 ≤ *k* ≤ *K*. Finally, we compute the eigenvectors as

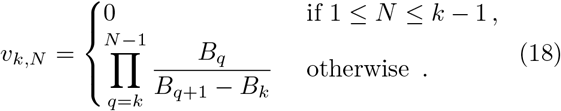

Notably, the eigenvectors **v**_*k*_ are ill-defined when the birth rates are degenerate (i.e., when *B*_*k*_ = *B*_*k*′_ for *k* ≠ *k*′). Indeed, equation (18) shows that some of the components of the eigenvectors diverge when birth rates degenerate. The degeneracy typically happens when the population birth rate curve displays particular symmetries (see figure 1). For instance, the Logistic growth model presents a mirror symmetry with respect to *K/*2; for all 1 ≤ *k* ≤ *K*, we obtain *B*_*k*_ = *B*_*K*−*k*_. Diagonalization of the transition rate matrix would thus not apply to the Logistic growth model.

In the case where every birth rate is distinct (i.e., *B*_*k*_ ≠ *B*_*k*′_ if *k* ≠ *k*′), we compute the constants *c*_*k*_ as

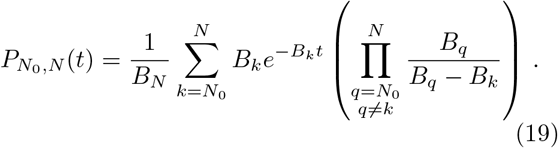

In the next section, we show how to extend this result to the cases where the birth rates may present some degeneracies.

### 4.2. Exact solution: distribution of waiting times between birth events

As we just saw, the method based on the transition rate matrix is unsatisfactory because it does not yield a closed-form solution for equation (14) if birth rates are degenerate. Here, we suggest a novel approach based on waiting times between birth events. The underlying idea is the following: in the absence of deaths, population growth can be interpreted as a succession of events (births) happening in a well-defined order separated by waiting times, which are random variables. To reach size *K* from its initial size *N*_0_, the population has to grow one individual at a time and go from *N*_0_ to *N*_0_ + 1, then from *N*_0_ + 1 to *N*_0_ + 2, etc. in this precise order. Therefore, all the information needed to derive the time-dependent probability 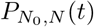 should be contained in the distribution of the time between two birth events. In other words, in the approach based on the master equation and the transition rate matrix, the reasoning is based on population sizes, whereas in this approach, our reasoning is instead based on waiting times between successive events.

We denote by *τ*_*N*_ the time elapsed between two births where the population size increases from *N* to *N* + 1 individuals. Owing to the Poissonian nature of the process, *τ*_*N*_ is a stochastic variable exponentially distributed with mean 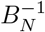. Then, the probability of having a given number *N* of individuals at time *t* must be equal to the probability that *N* − *N*_0_ births occurred by *t* and not *N* − *N*_0_ + 1 yet. Quantitatively speaking,

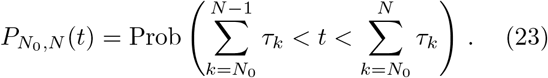

The sum of *n* exponentially distributed random variables with rates {*λ*_*i*_} _1≤*i*≤*n*_ follows a hypoexponential distribution [75, 76], which we denote *H* (*t*; {*λ*_*i*_}_1≤*i*≤*n*_). Hypoexponential distributions were previously studied in the context of population genetics [77] but also cell and systems biology [78–81].

Using the expression for the probability density function for the hypoexponential distribution (see table 3 for the three cases to consider), we write the exact solution to equation (7) in the form

**Table 3.**
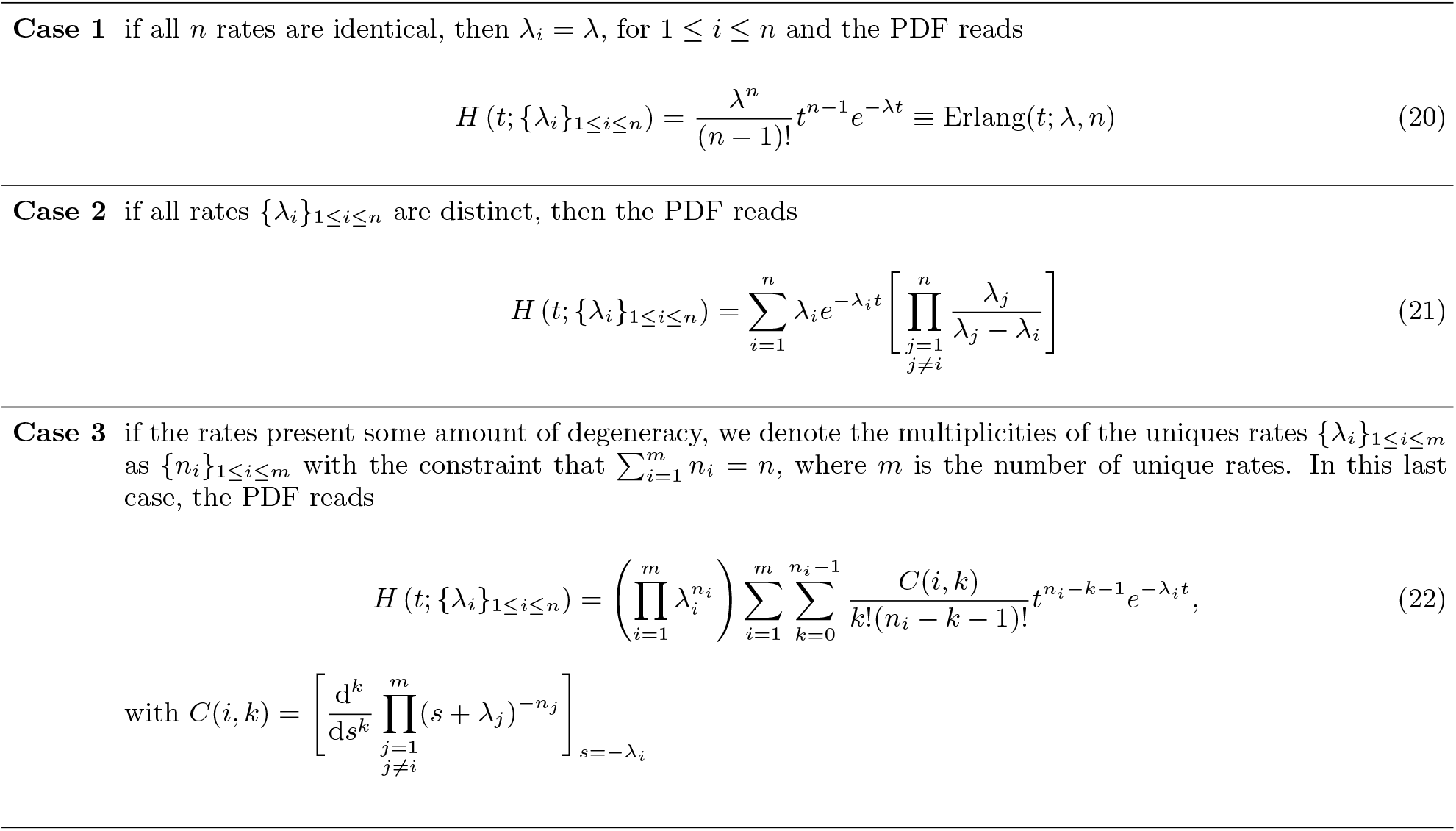
Definition of the probability density function for the hypoexponential distribution.

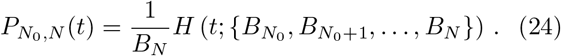

Notably, in the case where all population growth rates are distinct, the hypoexponential distribution takes the form (21), and we recover exactly the solution introduced in the previous section (see equation (19)). In figure 5, we compare our exact solution (24) to the mean population size measured in simulations of the stochastic process for our four nonlinear growth models and show perfect agreement in all cases. We also confirm in figure 5 that the full probability distributions measured from simulations agree with our exact solution.

## 5. APPLICATIONS

Finally, we cover two examples of applications in which we show that using a deterministic model instead of an exact solution to stochastic population growth leads to quantitatively very different results and may misinterpret important experimental results.

### 5.1. Population growth dynamics within a community

First, we extend our results to the study of a community composed of multiple strains evolving in the same environment. Community dynamics has received much attention recently with the growing field of microbiome studies, where predicting the relative abundance of each microbial strain in the gut microbiota represents an opportunity for medical diagnosis, and treatment [82]. For simplicity and without loss of generality, we focus on the case of two competing strains. Consider, for instance, the population dynamics of a wild-type (W) strain and a mutant strain (M) competing in a batch culture environment; we denote their intrinsic birth rates *b*_W_ and *b*_M_, respectively. As before, we define *N* as the size of the community, whereas *n* (resp. *N* − *n*) denotes the number of M (resp. W) individuals. We assume that the size of the community is limited by a single carrying capacity *K*. Note that our approach is easily generalizable to cases with multiple strains or with different carrying capacities and with explicit interaction parameters.

Furthermore, we introduce the relative fitness of the two strains, defined as *r* = *b*_M_*/b*_W_; this ratio indicates which strain is favored by natural selection. Specifically, if *r >* 1 (resp. *r <* 1), then strain *M* is beneficial (resp. deleterious), with *r* = 1 corresponding to the neutral case. Each time an individual reproduces, the probability that this individual is of strain M is then given by

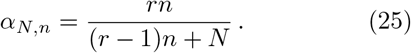

Here, we start with an initial community size *N*_0_, composed of *n*_0_ individuals from strain M and *N*_0_ −*n*_0_ individuals from strain W. The probability 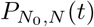 that the community has a *total* size *N* at time *t*, knowing that the initial size of the community was *N*_0_, is given directly by equation (24) with population reproduction rates

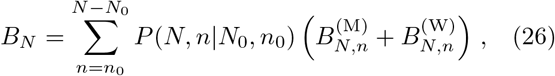

where 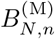 and 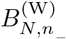 are the rates at which each population increases. For the Logistic model, these rates are for instance given by

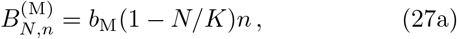

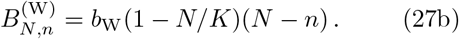

Furthermore, the probability *P* (*N, n*|*N*_0_, *n*_0_) of finding *n* individuals of type M when the total number of individuals is *N* satisfies

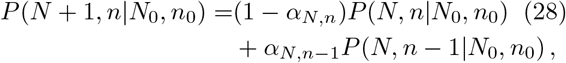

subject to the initial conditions 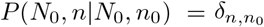. Note that equation (28) has been extensively studied in [83]. By definition, the probability to observe *n* individuals of type M at time *t*, knowing that we had initially *n*_0_ such individuals, is

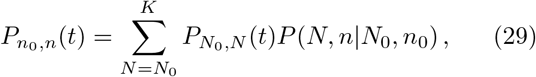

which we exactly compute using equations (24), and (28). Then, using equations (24), (26), and (28), we compute the average stochastic community and population sizes

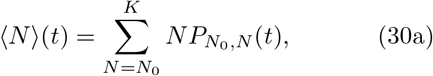

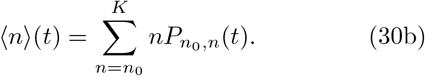

On the other hand, a deterministic description of the community dynamics leads to the system of differential equations

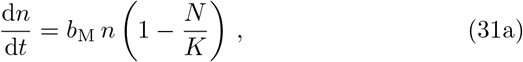

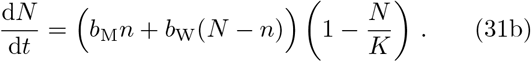

We once again compare the results of stochastic simulations quantitatively to the predictions of equation (30) on one hand and equation (31) on the other hand. We show in figure 6(a)-(b) that the deterministic model grossly overestimates the size of the community whereas our stochastic solution perfectly matches the simulated data. Strikingly, the deterministic model is shown to overestimate the equilibrium population size of strain M. Our stochastic approach provides an exact prediction of the average community and population sizes and, more importantly, yields the full time-dependent probability distributions of community and population sizes, which is not possible with a deterministic approach (see figure 6(c)-(d)).

**Figure 6.**
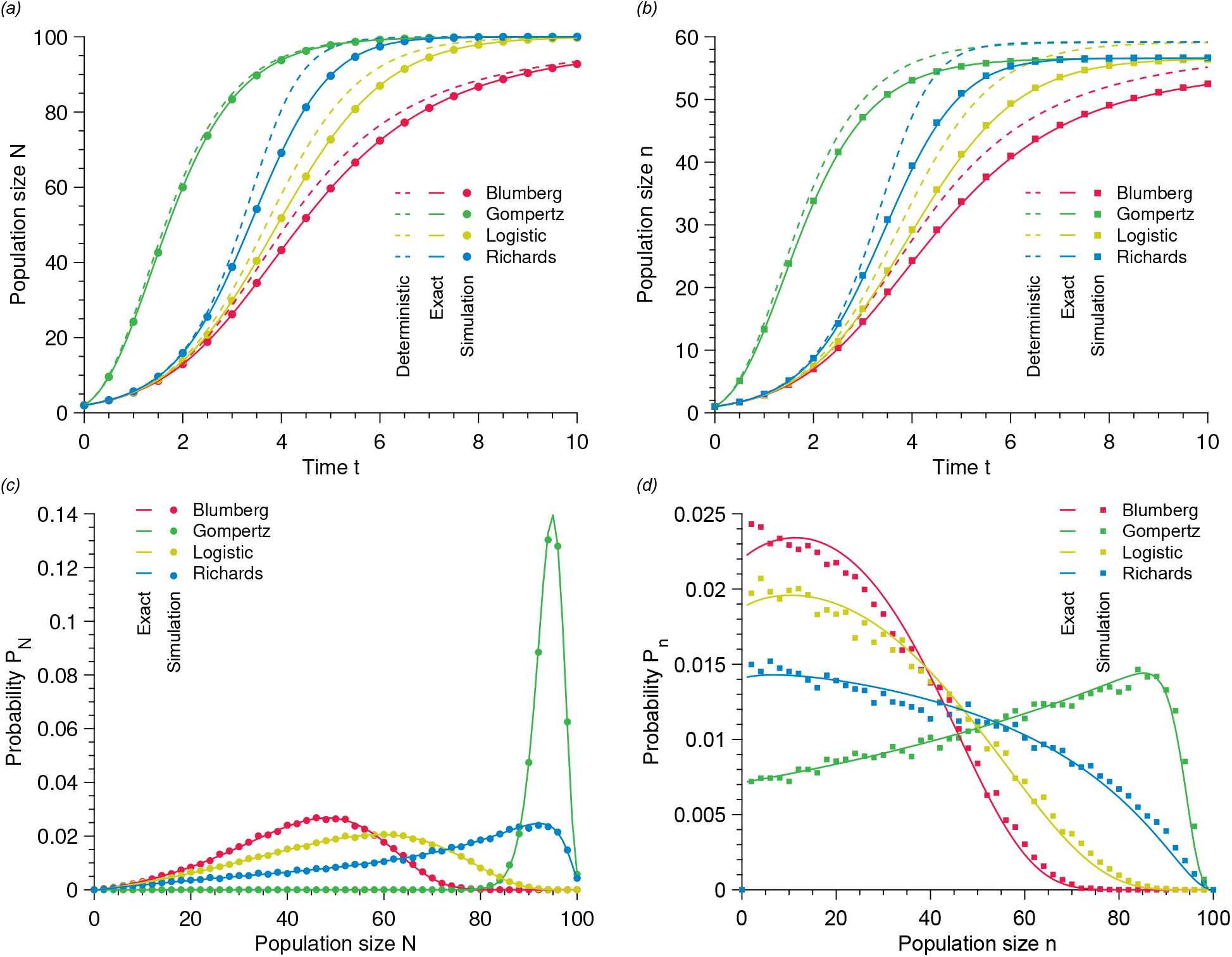
Exact time-dependent population growth and steady-state population sizes in community dynamics. (a) Total population size *N* versus time *t* for different population growth models. (b) Population size *n* of strain M versus time *t* for different population growth models. (c) Probability *P*_*N*_ of finding *N* individuals at time *t* = 4. Probability *P*_*n*_ of finding *n* individuals at time *t* = 4. In every panel, the solid lines represent our stochastic solution. In (a) and (b), the dashed lines show the deterministic predictions whereas each points results from simulated data averaged over 10^5^ stochastic realizations. Parameter values: carrying capacity *K* = 100, mutant birth rate *b*_*M*_ = 1.1, wild-type birth rate *b*_*W*_ = 1, initial community size *N*_0_ = 2 and initial wild-type population size *n*_0_ = 1.

### 5.2. Fixation probability in a serial passage experiment

Finally, we show that our exact stochastic solution yields the fixation probability of a strain in a serial passage experiment. For this, let us assume that we start an experiment with the same number of individuals M and W. As usual, the initial size of the community formed by strains M and W is given by *N*_0_; we introduce the dilution rate *D* defined as the ratio of the initial size of the community to the carrying capacity, *D* = *N*_0_*/K*.

In a serial passage experiment, the community grows for a time *τ* before one applies a bottleneck by taking a random sample of *N*_0_ individuals; mathematically, this corresponds to selecting *N*_0_ individuals from the community following a binomial law.

One then proceeds with a new growth phase of length *τ* before applying a new bottleneck. This process is repeated until only a single strain is left in the community. In these experiments, a quantity of interest is the probability *p*_fix_ that the strain M fixes and that the strain W goes extinct. In particular, optimizing the fixation probability as a function of the dilution ratio *D* and the waiting time *τ* has attracted a lot of attention, especially in the context of directed evolution [84–86]. To our knowledge, existing studies have only used deterministic models to answer this question.

To calculate *p*_fix_, we model the system as a Markov chain on the number of individuals M after each bottleneck event. At the moment of applying a bottleneck, the population contains *N* (*τ*) individuals, including *n*(*τ*) individuals of strain M. If picking a single individual randomly from the community, the probability that this individual is of type M is thus given by *n*(*τ*)*/N* (*τ*). The probability Π_*i*→*j*_ that the number of individuals M goes from *i* to *j* when a bottleneck is applied follows the binomial distribution

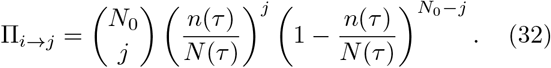

Under a deterministic approach, *n*(*τ*) and *N* (*τ*) are obtained by solving equation (31) with the initial conditions *n*_0_ = *i* and *N*_0_ = *DK*. However, in a stochastic approach, we write the Π_*i*→*j*_ as a sum over all possible pairs (*N, n*) at time *τ* weighted by their respective probabilities,

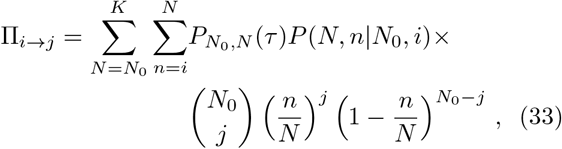

where *P* (*N, n*|*N*_0_, *n*_0_) is governed by equation (28). Finally, we note that conservation of probabilities imposes that Π_*j*→*j*_ = 1 −∑_*i* ≠*j*_ Π_*i*→*j*_.

We define 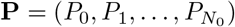 as the column vector of probabilities *P*_*i*_ to have *i* individuals of strain M in the random sample of *N*_0_ individuals from the community following a bottleneck event. Although the serial passage experiment defines a discrete-time Markov process, we follow reference [87] and take the limit of continuous time to write the master equation governing **P** as

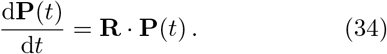

Here, the elements of the transition rate matrix **R** are given by

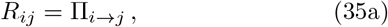

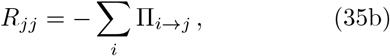

in which (35b) ensures conservation of probability.

The Markov process thus defined possesses two absorbing states, namely *n* = 0 and *n* = *N*_0_, which correspond to the extinction and fixation of strain M, respectively. By definition, once the system reaches one of these states, it remains there indefinitely. Mathematically, this implies that the first and last columns of the transition rate matrix are filled with zeros as these columns contain the transition rates out of the *n* = 0 and *n* = *N*_0_ states, respectively. Thus, the matrix **R** is not invertible. To deal with this issue, we introduce the reduced transition rate matrix 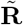 in which the rows and the columns corresponding to the absorbing states are removed. The fixation probability then reads

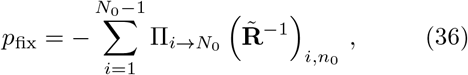

where *n*_0_ is the number of individuals M at the beginning of the experiment. As shown in figure 7, the deterministic approach grossly overestimates the fixation probability for all dilution rates *D*. Here, we confirm this result for all nonlinear growth models studied. On the other hand, we show that our stochastic solution exactly matches the results of stochastic simulations. This result has significant consequences as we show in particular that the dilution ratio predicted by a deterministic approach to optimize the fixation probability is far from being the *actual* optimal dilution ratio, although it is commonly used in the literature, as was already argued.

**Figure 7.**
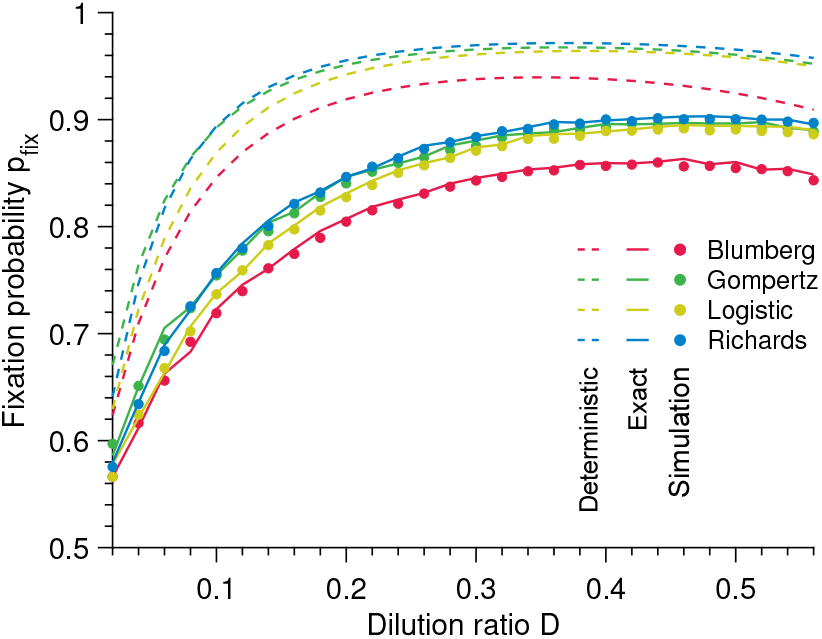
Fixation probability in a serial passage experiment. Fixation probability *p*_fix_ versus dilution ratio *D* for different population growth models. The solid lines represent our stochastic solution, the dashed lines the deterministic predictions, and each point results from simulated data averaged over 10^5^ stochastic realizations. Parameter values: carrying capacity *K* = 100, mutant birth rate *b*_M_ = 1.1, wild-type birth rate *b*_W_ = 1, time between each bottleneck *τ* = 3.

## 6. DISCUSSION

In conclusion, we showed that a deterministic approach to population growth leads to biased predictions of the average behavior of this inherently stochastic process. More precisely, deterministic models overestimate the population size averaged over large numbers of stochastic realizations, and this overestimation increases with decreasing initial population size. Qualitatively, the bias of the deterministic approach is due to the variability of the waiting times between reproduction events, which is particularly important at small population sizes. Quantitatively, the bias of the deterministic approach is due to unclosed-moment dynamics. Importantly, we showed that moment-closure approximations are not sufficient to significantly reduce the relative difference between analytical predictions and average population sizes, and are not applicable to all population growth models.

In contrast, we proposed two methods to derive exact solutions to the stochastic population growth dynamics: either by solving the master equation directly, which requires the diagonalization of the transition rate matrix, or by tracking reproduction times instead of population sizes. The first method was shown to be valid only in situations where the reproduction rates are distinct, whereas the second is generic. Our solution has revealed that the temporal distribution of population sizes is proportional to a hypoexponential distribution. Finally, we showed that our solution provides a more accurate description (than a deterministic approach) of the time-dependent and steady-state population sizes in a community composed of competing strains and the fixation probability of a mutation in a serial passage experiment.

Our theory offers an opportunity to quantify the dynamics of microbial communities from colonization to coexistence and thus contributes to the growing field of microbial eco-evolutionary dynamics. For example, the gut of *C. elegans* worms colonized by two neutrally-competing strains was shown to transition from a single-strain composition at a low colonization rate to coexistence at a high colonization rate [88]. In previous work, a deterministic approach was used to predict this transition [88]. Our exact solution enables the expansion of this work by quantifying the abundance distribution of either strain within each worm’s gut.

Our work opens new perspectives on population dynamics, ecology, and evolution. Importantly, our theory yields the full time-dependent probability distribution of population sizes [see equation (24)]. Based on this result, an interesting future research direction would be to improve inference methods for growth parameters, as current methods suffer from substantial limitations [49]. We propose that the present work constitutes the first step towards an exact inference method since it allows for the exact calculation of the likelihood function [89].

Here, we focused on the case where the only source of stochasticity in the problem resides in the waiting times between births, corresponding to a homogeneous population. A natural extension of our work would be to consider heterogeneous populations in which individuals display growth variability, i.e., different intrinsic birth rates *b*. In this context, it would be interesting to study the effect of this intrapopulation variability on the growth dynamics both computationally and analytically by extending the present work. Interestingly, a further source of stochasticity could be introduced in the initial population size by assuming that *N*_0_ is a random variable. This model assumption applies to the initial number of viruses or bacteria invading a host following infection, which is a stochastic variable.

## Supporting information

ESM

## Data availability

The authors state that all data necessary for confirming the conclusions presented in the article are represented fully within the article or Supplemental Material. Annotated C and Matlab implementations of numerical simulations are available at https://github.com/LcMrc.

## Authors’ Contributions

LM designed the study; LM and TB performed the numerical and analytical work; LM, CB and TB analyzed and interpreted the data; LM and TB wrote the manuscript; LM, CB and TB edited the manuscript.

## Acknowledgements

The authors thank the THEE Group and Stephan Peischl at UniBe for discussion and feedback on the manuscript.

