## Supplementary material for "Solving the stochastic dynamics of population growth": ESM

### Electronic Supplementary Material for "Solving the stochastic dynamics of population growth"

#### CONTENTS

1. Detailed simulation method 1
1.1. Single-species population 1
1.2. Community 1
1.3. Serial passage experiment 2

2. From the stochastic model to the deterministic limit 2
2.1. Diffusion approximation 3
2.2. Deterministic limit 3

3. Additional figures 4

References 4

#### 1. DETAILED SIMULATION METHOD

In this work, the population growths are simulated using a Gillespie algorithm.

##### 1.1. Single-species population

Let us denote by $N$ the population size. The single elementary event that can happen is the reproduction of an individual:

$$A \overset{b_N}{\to} 2A\,. \tag{1}$$

Simulations follow these elementary steps:

1. Initialization: The population starts from $N = N_0$ individuals at time $t = 0$.
2. The time increment $\Delta t$ is sampled randomly from an exponential distribution with mean $1/B_N$.
3. The population size increases from $N$ to $N + 1$ and the time from $t$ to $t + \Delta t$.
4. We go back to Step 2 and iterate until the total number of individuals is $K$ or the defined maximum time is reached.

##### 1.2. Community

Let us consider two types of individuals, namely M and W, whose numbers are denoted by $N_{\mathrm{M}}$ and $N_{\mathrm{W}}$, respectively. The two elementary events that can happen are the reproduction of an individual M or W:

$$M \overset{b^{(\mathrm{M})}_{N_{\mathrm{M}},N_{\mathrm{W}}}}{\to} 2M\,, \tag{2}$$

*

$$W \overset{b^{(\mathrm{W})}_{N_\mathrm{M},N_\mathrm{W}}}{\rightarrow} 2W\,. \tag{3}$$

Simulations follow these elementary steps:

1. Initialization: The population starts from $N_\mathrm{M} = N_{\mathrm{M},0}$ M individuals and $N_\mathrm{W} = N_{\mathrm{W},0}$ W individuals at time $t = 0$.

2. The time increment $\Delta t$ is sampled randomly from an exponential distribution with mean $1/\left(B^{(\mathrm{M})}_{N_\mathrm{M},N_\mathrm{W}} + B^{(\mathrm{W})}_{N_\mathrm{M},N_\mathrm{W}}\right)$. The next event that may occur is chosen randomly, proportionally to its probability. For instance, reproduction of a W individual is chosen with probability $B^{(\mathrm{W})}_{N_\mathrm{M},N_\mathrm{W}}/\left(B^{(\mathrm{M})}_{N_\mathrm{M},N_\mathrm{W}} + B^{(\mathrm{W})}_{N_\mathrm{M},N_\mathrm{W}}\right)$.

3. The time increases from $t$ to $t+\Delta t$ and the event chosen at Step 2 is executed. For instance, if reproduction of a W individual is chosen, then $N_\mathrm{W}$ increases by one.

4. We go back to Step 2 and iterate until the total number of individuals is $K$ or the defined maximum time is reached.

##### 1.3. Serial passage experiment

Let us consider two types of individuals, namely M and W, whose numbers are denoted by $N_\mathrm{M}$ and $N_\mathrm{W}$, respectively. The two elementary events that can happen are the reproduction of an individual M or W:

$$M \overset{b^{(\mathrm{M})}_{N_\mathrm{M},N_\mathrm{W}}}{\rightarrow} 2M\,, \tag{4}$$

$$W \overset{b^{(\mathrm{W})}_{N_\mathrm{M},N_\mathrm{W}}}{\rightarrow} 2W\,. \tag{5}$$

Simulations follow these elementary steps:

1. Initialization: The population starts from $N_\mathrm{M} = N_{\mathrm{M},0}$ M individuals and $N_\mathrm{W} = N_{\mathrm{W},0}$ W individuals at time $t = 0$. The next time when the dilution occurs is stored in the variable $t_\mathrm{dilution}$, which is initialized at $t_\mathrm{dilution} = \tau$, the first time when dilution occurs.

2. The time increment $\Delta t$ is sampled randomly from an exponential distribution with mean $1/(B^{(\mathrm{M})}_{N_\mathrm{M},N_\mathrm{W}} + B^{(\mathrm{W})}_{N_\mathrm{M},N_\mathrm{W}})$. The next event that may occur is chosen randomly, proportionally to its probability. For instance, reproduction of a W individual is chosen with probability $B^{(\mathrm{W})}_{N_\mathrm{M},N_\mathrm{W}}/(B^{(\mathrm{M})}_{N_\mathrm{M},N_\mathrm{W}} + B^{(\mathrm{W})}_{N_\mathrm{M},N_\mathrm{W}})$.

3. If $t + \Delta t < t_\mathrm{dilution}$, time is increased to $t + \Delta t$ and the event chosen at Step 2 is executed. For instance, if reproduction of a W individual is chosen, then $N_\mathrm{W}$ increases by one.

4. If $t + \Delta t \geq t_\mathrm{dilution}$, the event chosen at Step 2 is not executed, because a dilution has to occur before. The dilution is performed: time is incremented to $t = t_\mathrm{dilution}$, $N_{\mathrm{M},0}$ M individuals are selected from the community following a binomial law $\mathcal{B}(D \times K, N_\mathrm{M}/(N_\mathrm{M} + N_\mathrm{W}))$, and $N_{\mathrm{W},0} = D \times K - N_{\mathrm{M},0}$. In addition, $t_\mathrm{dilution}$ is incremented to $t_\mathrm{dilution} + \tau$, and thus stores the next time when the dilution occurs.

5. We go back to Step 2 and iterate until there is only one species left.

#### 2. FROM THE STOCHASTIC MODEL TO THE DETERMINISTIC LIMIT

Here, we present a full derivation of the deterministic limit of the stochastic pure-birth model. Starting from the recurrence equation satisfied by the probabilities of having a given population size, we obtain a Fokker-Planck equation, corresponding to the diffusion approximation, and then a deterministic differential equation, in the limits of increasingly large population sizes. Let us first recall the recurrence equation corresponding to the pure-birth process, where $N$ denotes the number of individuals:

$$P^{\tau+1}_{N_0,N} = B_{N-1}P^{\tau}_{N_0,N-1} + (1 - B_N)P^{\tau}_{N_0,N}\,. \tag{6}$$

Let us now introduce the reduced variables $n = N/K$, $t = \tau/N$, as well as $\rho(n,\tau|n_0) = K \times P^{\tau}_{N_0,N}$. Then, since one step of the pure-birth process occurs each time unit, Eq. 6 can be rewritten as:

$$\rho(n, t + 1/K|n_0) - \rho(n, t|n_0) = \rho(n - 1/K, t|n_0)B(n - 1/K) - \rho(n, t|n_0)B(n)\,. \tag{7}$$

##### 2.1. Diffusion approximation

For $K \ll 1$, considering that jumps are small at each step of the pure-birth process, i.e. $1/K \ll n$ and $1/K \ll t$, the probability density $\rho(n,t|n_0)$ and the transition probability $B(n)$ can be expanded in a Taylor series around $n$ and $t$. This expansion, known as a Kramers-Moyal expansion, yields, to first order in $1/K$:

$$\frac{\partial \rho(n,t|n_0)}{\partial t} = -\frac{\partial}{\partial n}\left(\rho(n,t|n_0)B(n)\right) + \frac{1}{2K}\frac{\partial^2}{\partial n^2}\left(\rho(n,t|n_0)B(n)\right) . \tag{8}$$

The previous equation is known as a diffusion equation, or a Fokker-Planck equation, or a Kolmogorov forward equation.

##### 2.2. Deterministic limit

In the limit $K \to \infty$, retaining only the zeroth-order terms in $1/K$, Eq. 8 reduces to:

$$\frac{\partial \rho(n,t|n_0)}{\partial t} = -\frac{\partial}{\partial n}\left(\rho(n,t|n_0)B(n)\right) . \tag{9}$$

Let us focus on the average value of $n$, denoted by $\langle n \rangle$. Using Eq. 9 yields

$$\frac{\mathrm{d}\langle n \rangle}{\mathrm{d}t} = \int_0^1 n \, \frac{\partial \rho(n,t|n_0)}{\partial t} \, \mathrm{d}n \, , \tag{10}$$

$$= -\int_0^1 n \, \frac{\partial}{\partial n}\left(\rho(n,t|n_0)B(n)\right) \mathrm{d}n \, , \tag{11}$$

$$= \left[n \, \rho(n,t|n_0)B(n)\right]_0^1 + \int_0^1 \rho(n,t|n_0)B(n)\mathrm{d}n \, , \tag{12}$$

$$= \langle B(n) \rangle \, . \tag{13}$$

The first term of right hand side of Eq. 12 vanishes because $B(0) = B(1) = 0$. In the limit $K \to \infty$, the distribution of $n$ is very peaked around its mean, so $\langle n \rangle \approx n$ and $\langle B(n) \rangle \approx B(n)$, yielding:

$$\frac{\mathrm{d}n}{\mathrm{d}t} = B(n) \, . \tag{14}$$

#### 3. ADDITIONAL FIGURES

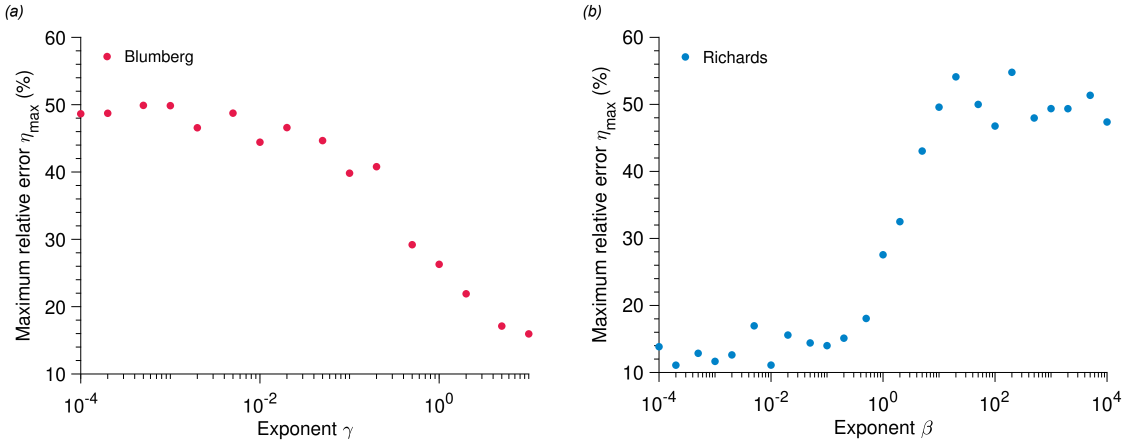


**Figure 1**. **The maximum relative error remains high for all parameter values.** Maximum relative error $\eta_{\max}$ versus exponent $\gamma$ (a) and $\beta$ (b) for the Blumberg and Richards models, respectively. Parameter values: $K = 100$, $N_0 = 1$, $b = 1$ and $\alpha = 1$.

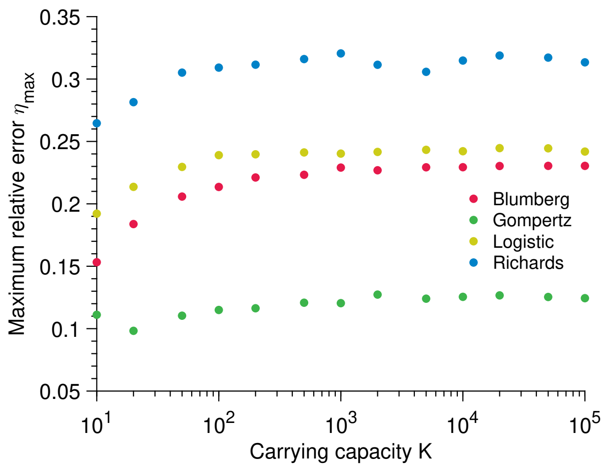


**Figure 2**. **The maximum relative error does not depend on the carrying capacity.** Maximum relative error $\eta_{\max}$ versus carrying capacity $K$ for different population growth models. Parameter values: $K = 100$, $N_0 = 1$, $b = 1$, $\alpha = 1$, $\beta = 2$ and $\gamma = 1.5$.
